# *Pyrenophora teres* and *Rhynchosporium secalis* infections in malt barley as influenced by genotype, spatial and temporal effects and nitrogen fertilization

**DOI:** 10.1101/649475

**Authors:** Petros Vahamidis, Angeliki Stefopoulou, Christina S. Lagogianni, Garyfalia Economou, Nicholas Dercas, Vassilis Kotoulas, Dionissios Kalivas, Dimitrios I. Tsitsigiannis

## Abstract

Net form net blotch (NFNB) and barley leaf scald are among the most important barley diseases worldwide and particularly in Greece. Their occurrence in malt barley can exert a significant negative effect on malt barley grain yield and quality. An experimental trial across two growing seasons was implemented in Greece in order i) to assess the epidemiology of NFNB and leaf scald in a barley disease free area when the initial inoculation of field occurs through infected seeds, and ii) to further explore the relationship among nitrogen rate, grain yield, quality variables (i.e. grain protein content and grain size) and disease severity and epidemiology. It was demonstrated that both NFNB and leaf scald can be carried over from one season to the next on infected seed under Mediterranean conditions. However, disease severity was more pronounced after barley tillering phase when soil had been successfully inoculated first. When nitrogen rate and genotype were the main sources of variation the epidemiology assessment was implemented with hotspot and Anselin Local Moran’s I analysis. It was found that the location of hotspots was modified during growing season. Soil and plant variables were assessed for the explanation of this variability. According to commonality analysis the effect of distance from the locations with the highest disease infections was a better predictor of disease severity (for both diseases) compared to nitrogen rate during pre-anthesis period. However, disease severity after anthesis was best explained by nitrogen rate only for the most susceptible cultivars to NFNB. The effect of disease infections on yield, grain size and grain protein content varied in relation to genotype, pathogen and stage of crop development. The importance of crop residues on the evolution of both diseases was also highlighted.

## Introduction

Barley (*Hordeum vulgare* L) is one of the leading cereal crops of the world and it is clearly number two in Europe in terms of cultivated acreage, next to bread wheat (*Triticum aestivum* L.) [1]. According to Meussdoerffer and Zarnkow [2], barley is the major source for brewing malts, which constitute the single most important raw material for beer production. *Pyrenophora teres* f. *teres* an ascomycete that causes the foliar disease net form net blotch (NFNB) and *Rhynchosporium secalis,* causal agent of barley leaf scald are among the most important barley diseases worldwide [3-5]. It is estimated that both these diseases can decrease barley grain yield up to 30–40% [3, 5-11]. In addition, there are indications that these diseases can also have a negative effect on malt barley quality [5].

Only a few studies have addressed so far the impact of NFNB and leaf scald on malt barley quality [12-13], and their results were restricted to northern climates. However, there is a lack of evidence on what really happens under Mediterranean conditions where the occurrence of malt barley diseases coincides with terminal drought. Malt barley has to meet certain specific quality requirements according to malt industry demands. Grain size and grain protein content are among the most important quality factors for malting barley [14]. Although the average grain weight and size is primarily determined during the post-anthesis period [15-16], grain protein content can also be affected during the pre-anthesis period. For example, pre-anthesis drought stress can cause a low nitrogen uptake during the vegetative period, thus reducing the yield potential. Then, more nitrogen is available during grain filling due to the low number of seeds, and grain protein content is increased [17].

Nitrogen fertilizer rate plays a major role in malt barley by affecting to a great extent the final yields, grain protein content (that has to be maintained below a threshold of 11.5-12.0% depending on brewing industry), as well as the susceptibility to leaf diseases. More nitrogen can increase the yield of malt barley [18-21], but can also exert an adverse effect on quality by increasing grain protein content [14, 22-24]. In addition, high nitrogen rates can also increase the susceptibility of barley to leaf diseases [13, 25-28]. Therefore, understanding the relationship among nitrogen rate, grain yield, quality variables and leaf disease infections can be very useful to further raising yield and to maintain the quality at a level that meets the requirements of malt industry.

In this study we aimed, i) to estimate the epidemiology of NFNB and leaf scald in a barley disease free area when the initial inoculation of the field occurs through infected seeds, and ii) to further explore the relationship among nitrogen rate, grain yield, quality variables (i.e. grain protein content and grain size) and disease severity and epidemiology.

## Materials and methods

### Study site and experimental design

The experiment was divided into three different phases, namely: a) the selection of malt barley seeds from infected crops (i.e. NFNB and leaf scald) grown in the main productive areas of malt barley in Greece (growing season 2013-2014), b) the inoculation year (Exp 1; growing season 2014-2015) when the seeds from the infected malt barley cultivars (i.e. *cv*. Grace, *cv*. Charles, *cv*. Fortuna, *cv*. KWS Asta and *cv*. Zhana) were grown in a barley disease free area and c) the application in the same location (i.e. inoculated soil with infected crop residues from Exp1) of nitrogen treatments on the most important (in terms of harvested areas) malt barley cultivars in Greece namely *cv*. Zhana, *cv*. Grace, *cv*. Traveler and *cv*. RGT Planet (Exp 2; growing season 2015-2016).

The experiments (Exp1 and Exp2) were conducted in Spata, Greece (37°58’44.34”N, 23°54’47.87”E and 118 m above sea level), at the experimental station of the Agricultural University of Athens. The soil was clay loam. Physical and chemical characteristics of the soil at the beginning of the experiments (November 2013) were: pH 7.7 (1:1 soil/water extract), organic matter 2.02%, CaCO_3_ 27.80%, electrical conductivity (*Ec*) 0.29 mmhos cm^-1^, total N (Kjeldahl) 0.105%, available P (Olsen) 52.84 ppm and 452 ppm exchangeable K.

In Exp1 the treatments consisted of 5 five malt barley cultivars as stated above. The experimental design was a randomized complete block design with 9 replications (in order to have a better spatial distribution of the selected genotypes) per genotype. During the second year (Exp2) the experiment was arranged in a two factorial randomized complete block design with three replications. Treatments were completely randomized within each block and included four two-rowed malt barley (*H. vulgare* L.) cultivars (i.e. *cv*. Zhana, *cv*. Grace, *cv*. Traveler and *cv*. RGT Planet) and four nitrogen fertilization rates. The four N application rates were 0 (N0), 60 (N1), 100 (N2) and 140 (N3) kg N ha^-1^. In order to achieve a more efficient use of the N, half of it was applied to the experimental plots at the onset of tillering phase (stage 20-22 according to Zadoks et al., 1974 scale) and the remaining at the end of tillering phase (stage 25-29 according to Zadoks scale [29]) as ammonium nitrate.

In both experimental years plot size was 9 m^2^ including 15 rows with row space of 20 cm and the crops were planted at a seed rate of approximately 350 seeds m^-2^. The plots in Exp2 were established in the same location where the plots of Exp1 had been seeded. In Exp1 sowing was carried out following conventional soil tillage (i.e. ploughing and then disc cultivator), whereas only rotary cultivator was used in Exp2 in order to simulate conditions of increased soil-borne disease pressure. Only certified malt barley seeds were used in Exp2, therefore the only source of disease dispersal was the crop residues from Exp1.

Soil water content was frequently determined during each cultivation season. EC-5 sensors of Decagon Devices, Inc. were installed at 25 cm depth in four different plots for the monitoring of the soil water content (SWC).

### Disease assessment

A slight modification of the equation proposed by Saari and Prescott [30] was adopted to estimate the disease severity (DS) during the phenological stages of tillering, stem elongation and milk development:

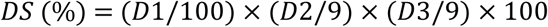

Where *D1* is the percentage of diseased plants in each plot, *D2* is the height of infection (i.e. 1=lowest leaf; 2=second leaf from base; 3-4=second leaf up to below middle plant; 5=up to middle of plant; 6-8= from center of plant to below the flag leaf; 9=up to flag leaf) and *D3* is the extent of leaf area affected by disease (i.e. 1=10% coverage to 9 = 90% coverage).

The area under disease progress curve (AUDPC) was calculated by following the formula given by Shaner and Finney [31]:

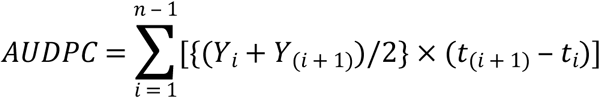

Where, *Y*_*i*_ = disease level at time *t*_*i*_ (*t*_*(i+1)*_ - *t*_*i*_) is the interval between two consecutive assessments and *n* is the total number of assessments.

Barley varieties were naturally infected by both diseases. The pathogens were further identified in the lab [4].

### Yield and malt characters measurements

At maturity, grain yield estimation was based on an area of 1 m^2^ per plot. Grain size was determined by size fractionation using a Sortimat (Pfeuffer GmbH, Kitzingen, Germany) machine, according to the 3.11.1 Analytica EBC “Sieving Test for Barley” method [32]. Nitrogen content was determined by the Kjeldhal method and protein content was calculated by multiplying the N content by a factor of 6.25 as described by Vahamidis et al. [33].

### Spatial statistical analysis

Using the geographical coordinates of the experimental plots, ArcGIS 10 was used to explore spatial associations, based on autocorrelation indices, of the disease severity among the experimental plots during the different developmental stages. Global autocorrelation indices, like Moran’s I, assess the overall pattern of the data and sometimes fail to examine pattern at a more local scale [34]. Thus, aiming at deepening our knowledge on spatial associations, local autocorrelation indices were used to compare local to global conditions. In this framework, hotspot analysis was used to identify statistically significant clusters of high values (hot spots) and low values (cold spots) using the Getis-Ord Gi statistic. Anselin Local Moran’s I was used to identify spatial clusters with attribute values similar in magnitude and specify spatial outliers.

In order to further explore the relationship between crop residues and disease severity, the distance between the crop residues of the previous season (2014/2015) and the location of the experimental plots of the investigated growing season (2015/2016) was calculated. At this point, it should be mentioned that Zhana was the only cultivar that was infected by *Rhynchosporium secalis* and Grace was the cultivar with the highest infection by *Pyrenophora teres* f. *teres*.

### Hotspot analysis

Moran’s I is a popular index to globally assess spatial autocorrelation, however it does not efficiently recognize the grouping of spatial patterns [35]. Hotspot analysis was used to assess whether experimental plots with either high or low values cluster spatially. Hotspot analysis uses the Getis-Ord local statistic given as:

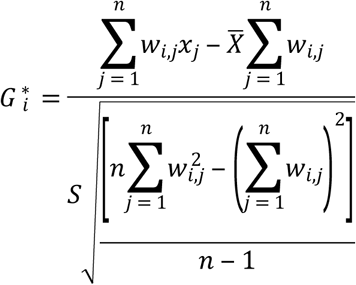

Where x_j_ is the disease severity value for experimental plot j, w_i,j_ is the spatial weight between experimental plot i and j, n is the total number of experimental plots and

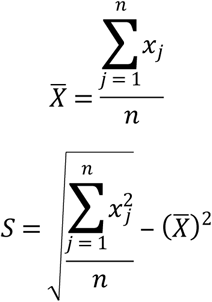

The Getis -Ord Gi statistic assesses whether the neighborhood of each experimental plot is significantly different from the study area and can distinguish high values clusters (hot spots) and low values clusters (cold spots).

The Gi* statistic returns a z-score which is a standard deviation. For statistically significant positive z-scores, higher values of z-score indicate clustering of high values (hot spot). For statistically significant negative z-scores, lower values indicate clustering of low values (cold spot).

### Cluster and outlier analysis

Anselin Local Moran’s I was used to identify clusters and spatial outliers. The index identifies statistically significant (95%, p<0.05) clusters of high or low disease severity and outliers. A high positive local Moran’s I value implies that the experimental plot under study has similarly high or low values as its neighbors, thus the locations are spatial clusters. Spatial clusters include high–high clusters (high values in a high value neighborhood) and low–low clusters (low values in a low value neighborhood). A high negative local Moran’s I value means that the experimental plot under study is a spatial outlier [36]. Spatial outliers are those values that are obviously different from the values of their surrounding locations [37]. Anselin Local Moran’s I enables us to distinguish outliers within hot spots, because it excludes the value of the experimental plot under study, in contrary to the hotspot analysis, which takes it into account.

Local Moran’s I is given as:

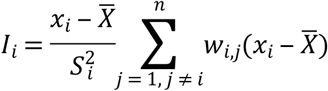

Where x_i_ is an attribute for feature I, 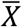 is the mean of the corresponding attribute, w_i,j_ is the spatial weight between feature I and j, and:

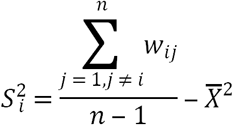

### Statistical analysis

Analyses of variance was performed using Statgraphics Centurion ver. XVI software package (Statpoint Technologies, Inc.,USA, Warrenton, Virginia). The experiment was a 2 × 4 factorial, replicated three times in a randomized complete block design. Significant differences between treatment means were compared by the protected least significant difference (LSD) procedure at *P* < 0.05. Commonality analysis was performed in the R Environment (version 3.4.3) using the ‘yhat’ package (version 2.0-0) as described by Nimon et al. [38].

## Results

### Weather conditions

The weather regime, in terms of maximum (T*max*) and minimum air temperature (T*min*) and rainfall, during both experiments is presented in Fig 1. The maximum and minimum temperatures increased from February to May, as typically occurs in Mediterranean environments. Environmental conditions differed between the two experimental years, with differences in the amount and distribution of precipitation during the growing season, as well as in temperature. In general 2015-2016 (Exp2) was considered to be a dryer growing season compared to 2014-2015 (Exp1).

**Fig 1.**
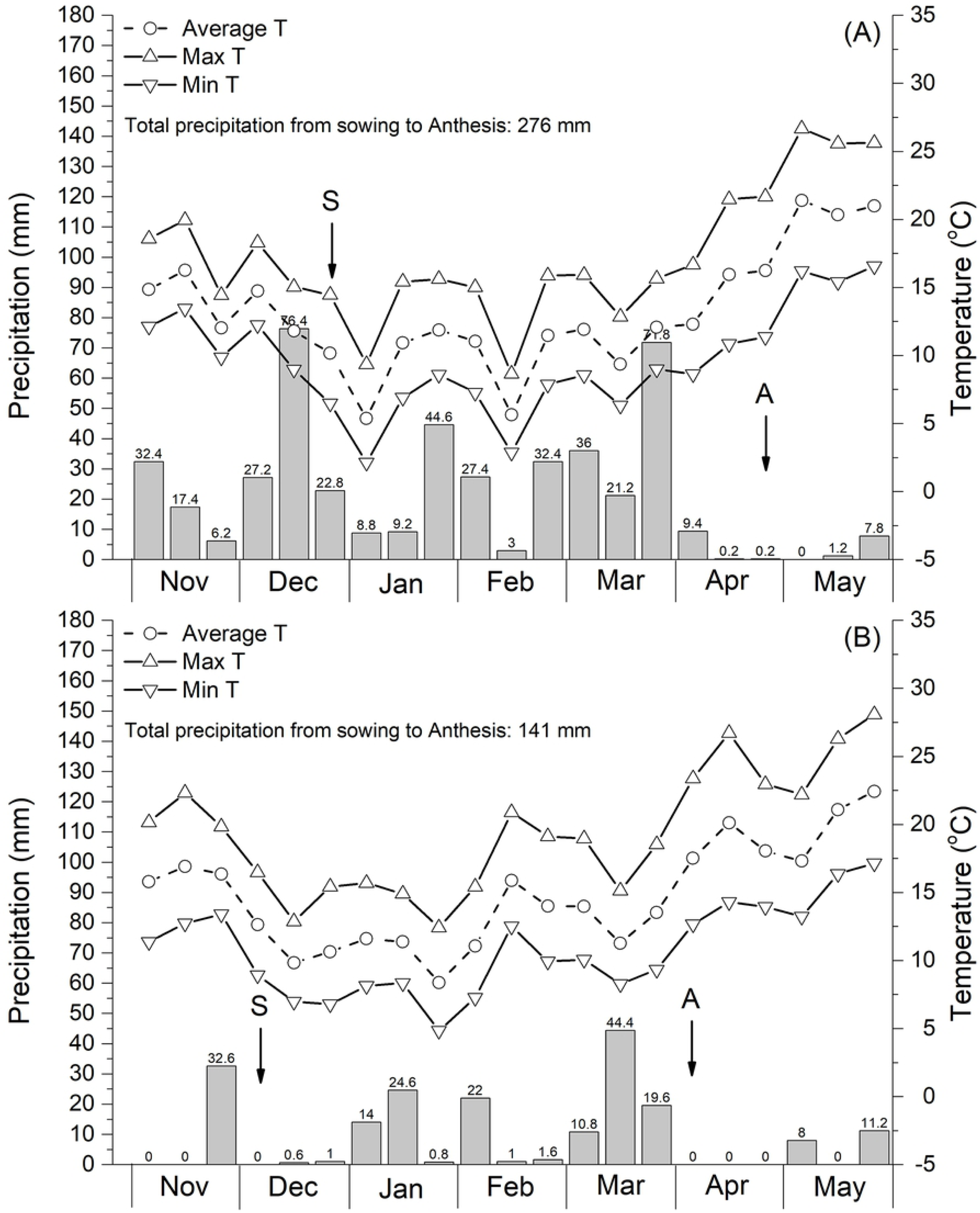
Precipitation and air temperature (*T*min and *T*max) during Exp1 (A, 2014-2015) and Exp2 (B, 2015-2016). The arrows indicate the main phenological stages: S=sowing; A=Anthesis.

### Temporal and genotypic effects

Charles, Grace, Traveler, Fortuna, KWS Asta and RGT Planet were exclusively infected with *Pyrenophora teres f. teres* (net form net blotch - NFNB), whereas the cultivar Zhana was exclusively infected with *Rhynchosporium secalis* (leaf scald). NFNB occurred at all developmental stages and in both experiments, whereas leaf scald was consistently observed after the onset of stem elongation phase (Fig 2). Although disease severity tended to be higher in Exp1 (disease dispersal from infected barley seed) compared to Exp2 (diseases dispersal from infected barley debris left after harvest) during the tillering phase of malt barley, after the onset of stem elongation stage it was more pronounced in Exp2. The same trend was also observed with leaf scald. In general, infections by NFNB were more severe compared to those by leaf scald, during all tested developmental phases of malt barley (Fig 2).

**Fig 2.**
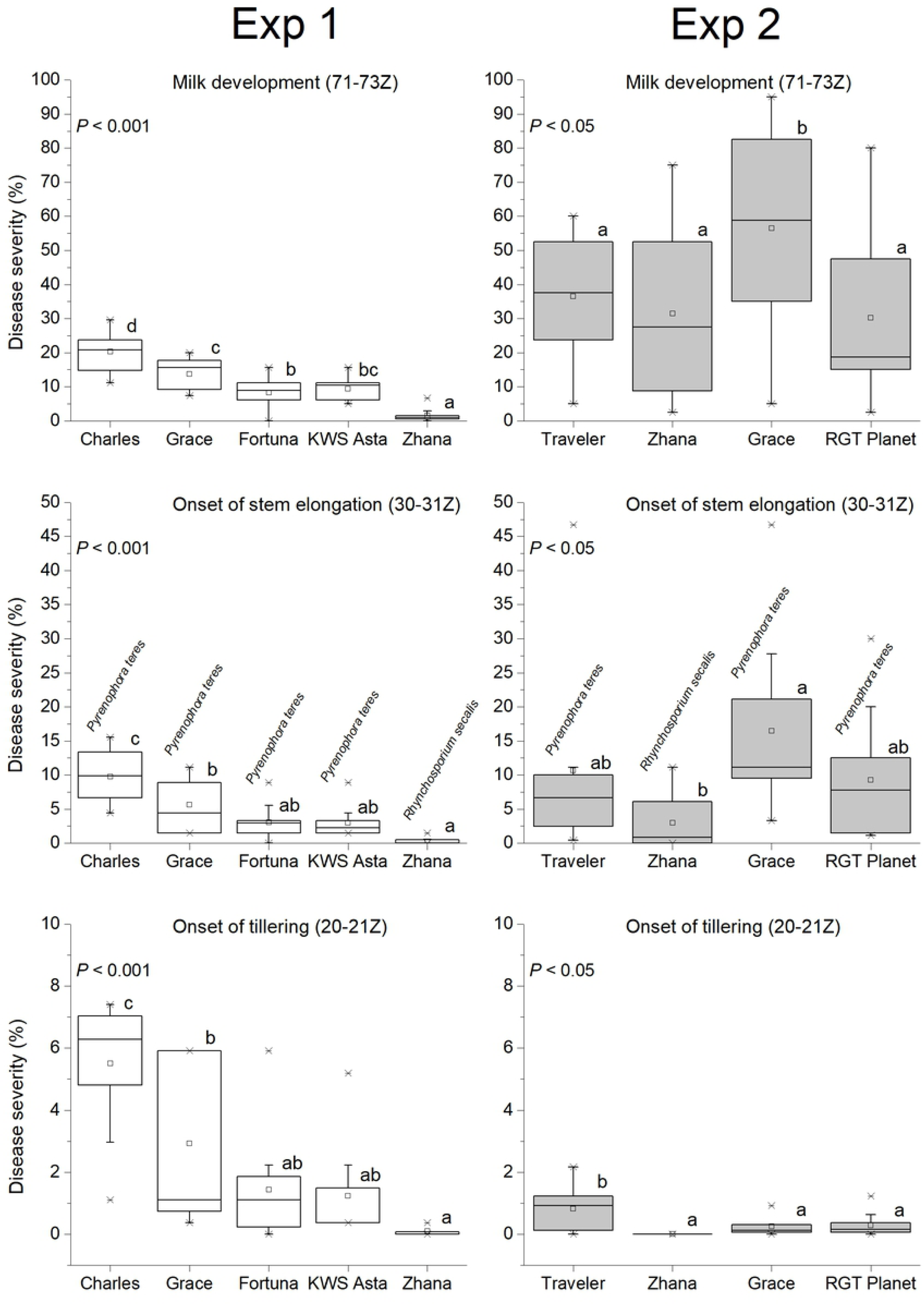
Malt barley cultivars susceptibility to *Pyrenophora teres f. teres* (net form net blotch - NFNB) and *Rhynchosporium secalis* (leaf blotch, scald) at different developmental phases during both experiments. The numbers in the brackets refer to Zadoks scale. Broad lines are medians, square open dots are means, boxes show the interquartile range and whiskers extend to the last data point within 1.5 times the inter-quartile range. *P*-values of ANOVA and permutation tests are given. Groups not sharing the same letter are significantly different according to L.S.D. test (*p* < 0.05).

**Fig 3.**
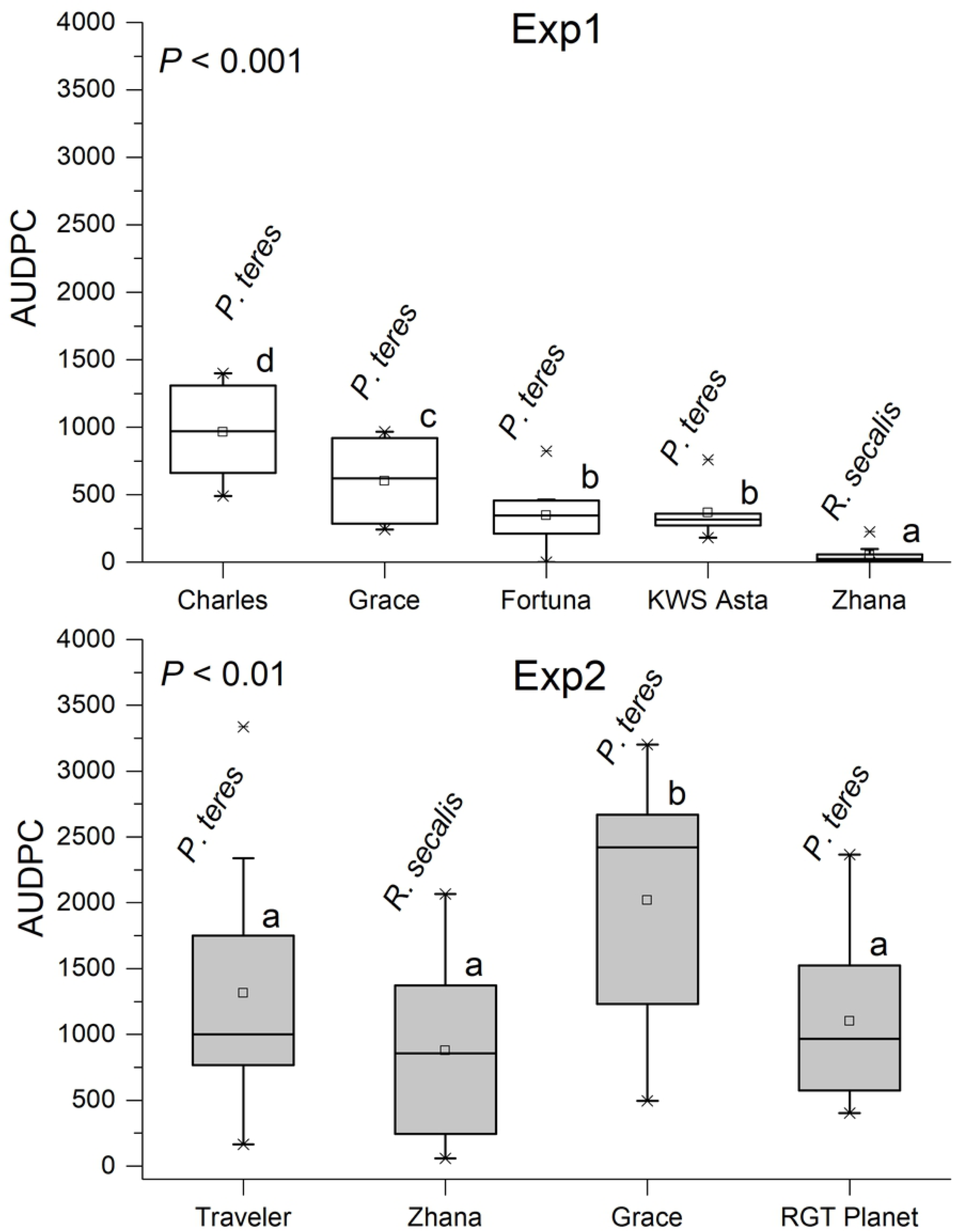
Malt barley cultivars susceptibility to *Pyrenophora teres f. teres* (net form net blotch - NFNB) and *Rhynchosporium secalis* (leaf blotch, scald) based on the area under disease progress curve (AUDPC). Broad lines are medians, square open dots are means, boxes show the interquartile range and whiskers extend to the last data point within 1.5 times the inter-quartile range. P-values of ANOVA and permutation tests are given. Groups not sharing the same letter are significantly different according to L.S.D. test (*p* < 0.05).

### Effect of N and genotype on grain yield and quality characters

Although the experimental data demonstrated a tendency for a positive relationship between the disease severity during grain filling and the rate of applied nitrogen (Fig 4), this tendency was not expressed in a statistical significant way according to ANOVA (Table 1). The only variable that was significantly affected by the rate of applied nitrogen was grain protein content (Table 1). With the exception of Zhana (i.e. it was the only cultivar that was infected with *Rhynchosporium secalis*) an increased disease severity generally resulted in higher grain protein content. However, it was recorded a genotypic variation among the studied cultivars concerning their response to increased disease severity (Fig 5).

**Table 1.** ANOVA summary for grain yield, grain protein content, maltable (% grains > 2.2 mm), AUDPC and disease severity (DS) during the onset of stem elongation and grain filling phases

| Source of variation | Grain yield (kg/ha) | Grain protein content (%) | Maltable (%) | AUDPC<sup>a</sup> | DS (stem elongation) | DS (grain filling) |
| --- | --- | --- | --- | --- | --- | --- |
| Cultivar | ** | ns | *** | ** | * | * |
| Nitrogen | ns | *** | ns | ns | ns | ns |
| Cultivar x Nitrogen | * | ns | ns | ns | ns | ns |

\*, \*\*, \*\*\* F values significant at the $P < 0.05$ , $P < 0.01$ and $P < 0.001$ probability levels, respectively.
ns stands for non-significant effect.
<sup>a</sup>AUDPC: Area under disease progress curve.

**Fig 4.**
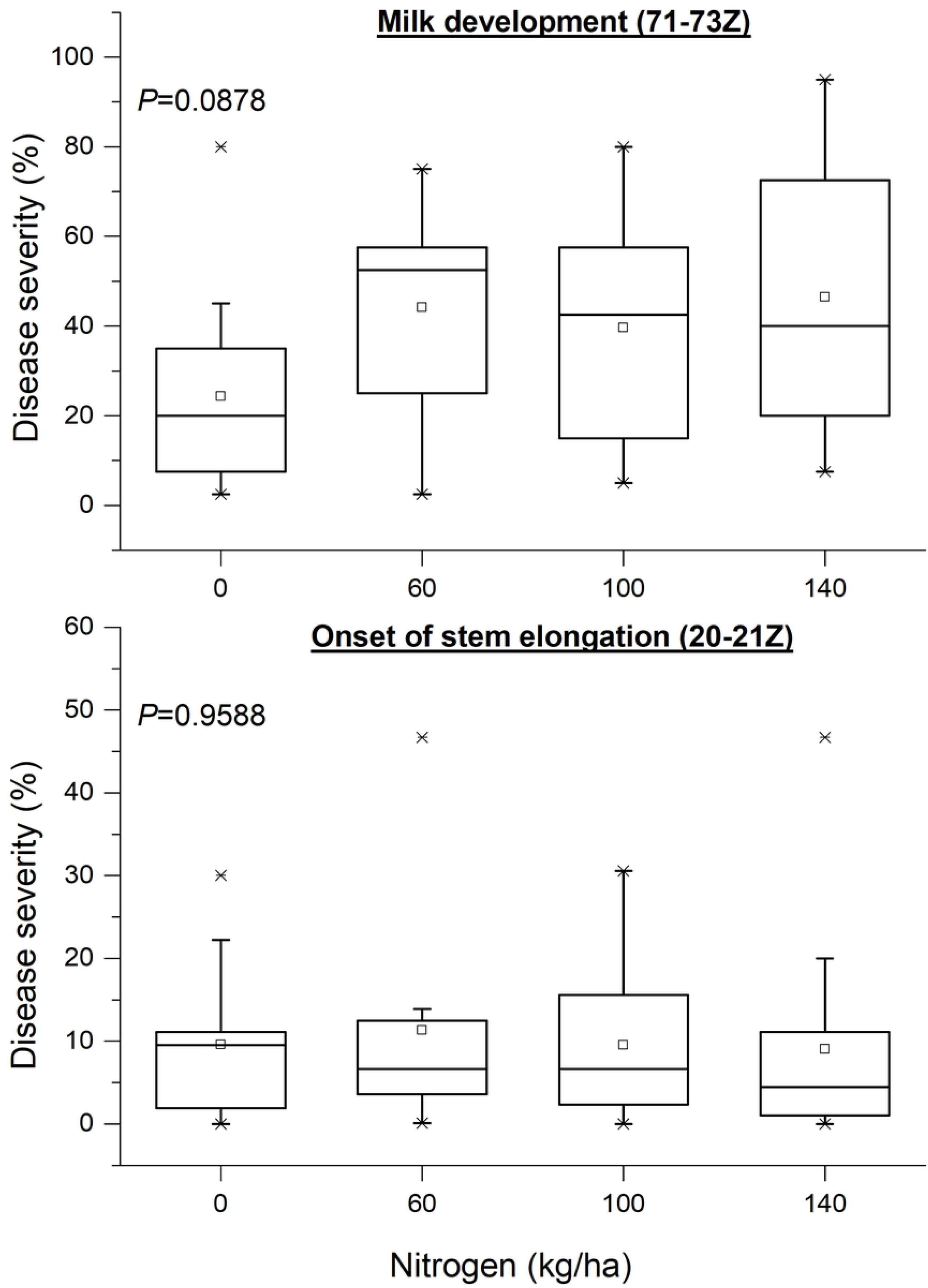
The effect of nitrogen rate on disease severity (caused by *Pyrenophora teres f. teres* and *Rhynchosporium secalis*) assessed at different developmental stages of malt barley. Broad lines are medians, square open dots are means, boxes show the interquartile range and whiskers extend to the last data point within 1.5 times the inter-quartile range. P-values of ANOVA and permutation tests are given.

**Fig 5.**
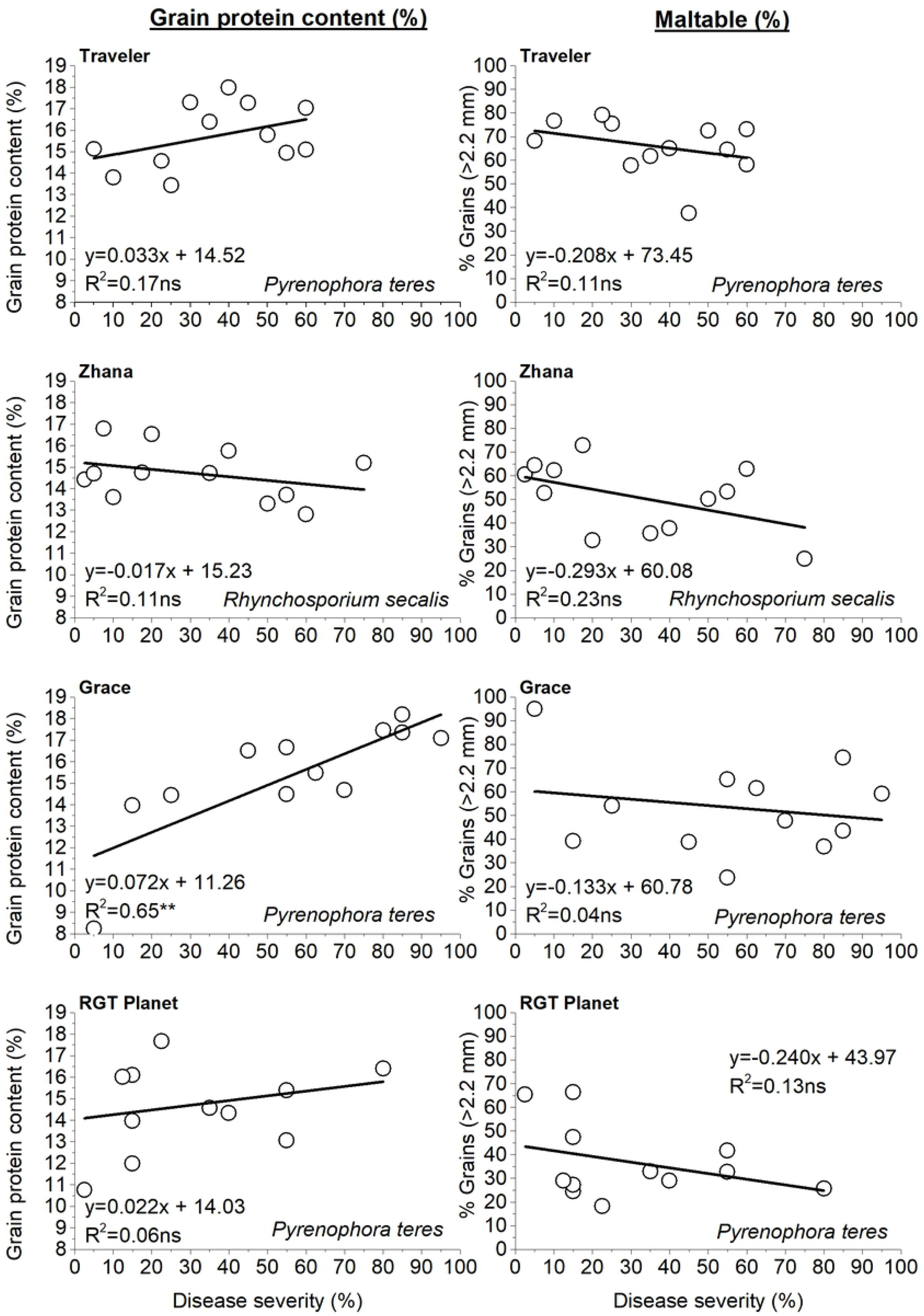
Relationship among disease severity (caused by *Pyrenophora teres f. teres* and *Rhynchosporium secalis*) with grain protein content and maltable grain size fraction (>2.2 mm) at grain filling phase when the main source of variation is nitrogen rate. *At *P*≤ 0.05; **At *P*≤ 0.01; ***At *P*≤ 0.001; ns=non-significant.

Grain yield was significantly affected by cultivar and by the interaction cultivar x nitrogen (Table 1), and varied from 0.84 to 4.26 t ha^-1^. Grace and Traveler were the only cultivars that presented significant relationships between grain yield and disease severity (Fig 6). In particular, Traveler recorded a marginal statistically significant negative relationship between grain yield and disease severity, only for the period of tillering (Fig 6). Concerning Grace, grain yield showed a negative significant direct relationship to disease severity for the period of grain filling (milk development) and on the contrary, presented a moderate positive association to disease severity for the period of tillering phase (Fig 6).

**Fig 6.**
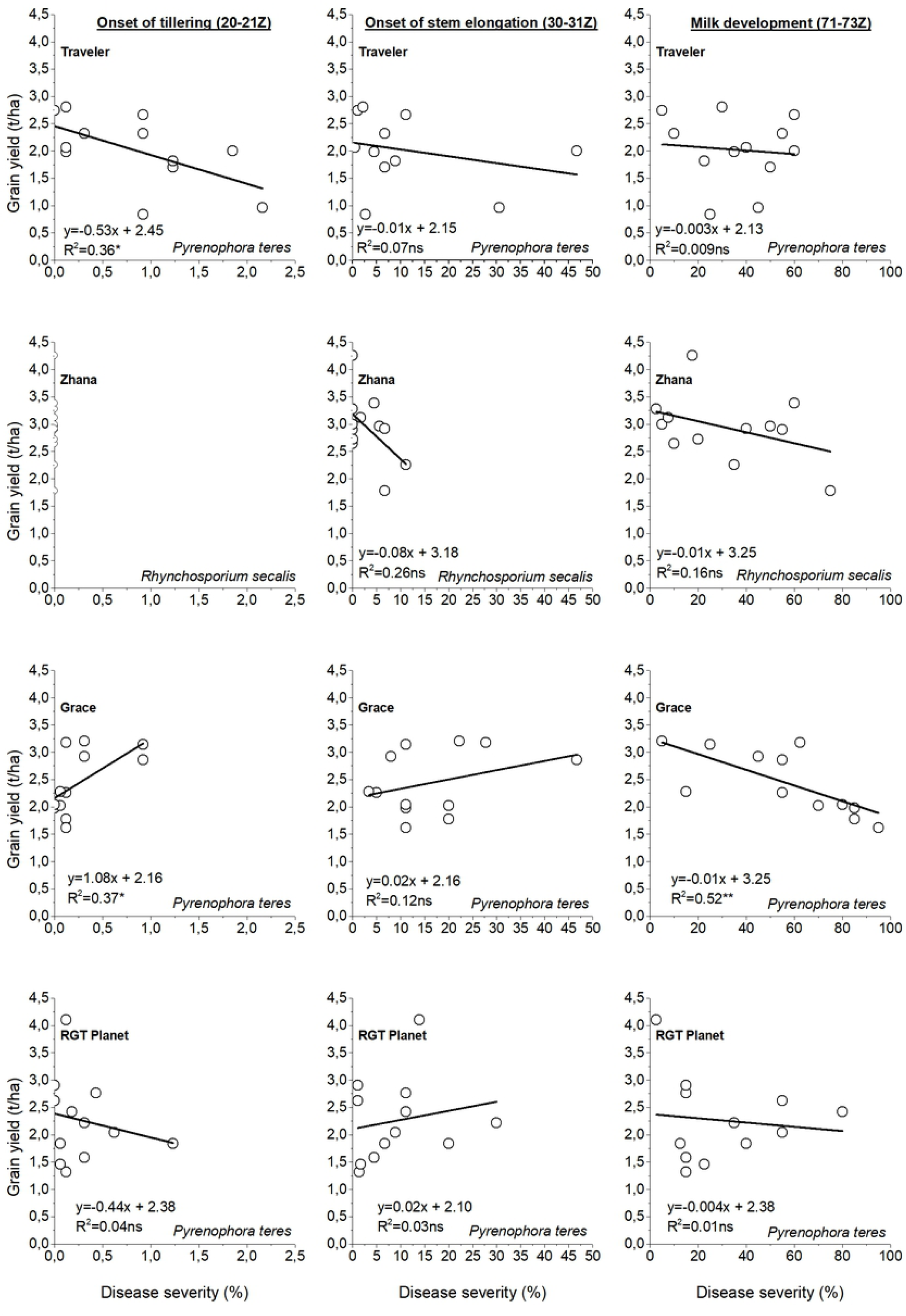
Relationship between grain yield and disease severity (caused by *Pyrenophora teres f. teres* and *Rhynchosporium secalis*) assessed at different developmental stages of malt barley, when the main source of variation is nitrogen rate. The numbers in the brackets refer to Zadoks scale. *At *P*≤ 0.05; **At *P*≤ 0.01; ***At *P*≤ 0.001; ns=non-significant.

The proportion of maltable grain size fraction (% grains > 2.2 mm), as well as disease severity during stem elongation and grain filling phases were not significantly affected by the rate of applied nitrogen (Table 1). A negative, but not significant, association was recorded between the proportion of maltable grain size fraction and disease severity for all the studied cultivars (Fig 5).

### The area under disease progress curve (AUDPC)

The area under disease progress curve (AUDPC) in Exp2 was not significantly affected either by nitrogen rate or the interaction cultivar x nitrogen (Table 1). However, the analysis of variance for AUDPC indicated that a significant degree of genotypic variation existed among the studied malt barley cultivars in both experiments. The AUDPC values were lower in Exp1 compared to Exp2. Charles and Grace presented the highest values in Exp1 and Exp2, respectively (Fig 3).

### Epidemiology assessment when nitrogen rate and genotype are the main sources of variation

Distribution patterns of disease severity were analyzed by using hotspot and cluster and outlier analysis in ArcGIS 10x for three different crop developmental periods: 1) tillering (20-21Z), 2) stem elongation (30-31Z) and 3) milk development (71-73Z). Cluster and outlier analysis was used to identify clusters of disease infected areas in cluster types of HH, HL, LL, and LH. LH represents a cluster of low values surrounded by high values, while HL is a cluster of high values surrounded by low values. In addition, LL and HH were statistically significant (p < 0.05) clusters of low and high disease severity values, respectively.

During the onset of tillering phase, two experimental plots presented significant positive z scores demonstrating significant clusters of intense disease severity. They were located on the western part of the field and both of them included Traveler with nitrogen rate of 100 and 140 kg/ha, respectively (Fig 7). A further investigation revealed that the distance of Traveler experimental plots from the previous season crop residues (i.e. the sites with Grace) explained 34% of the variation in disease severity (Fig 8). RGT Planet with nitrogen rate of 100 kg/ha, was also marked as a hotspot, but less intense since it presents a lower z score (Fig 7). It is reminded that lower z-scores indicate less intense clustering. The Local Moran’s I spatial analysis, indicated only one High-Low outlier in the western part of the field. Indeed, Traveler with a rate of 100 kg N /ha was considered as an outlier since it presented high values of disease severity surrounded by lower surrounding values.

**Fig 7.**
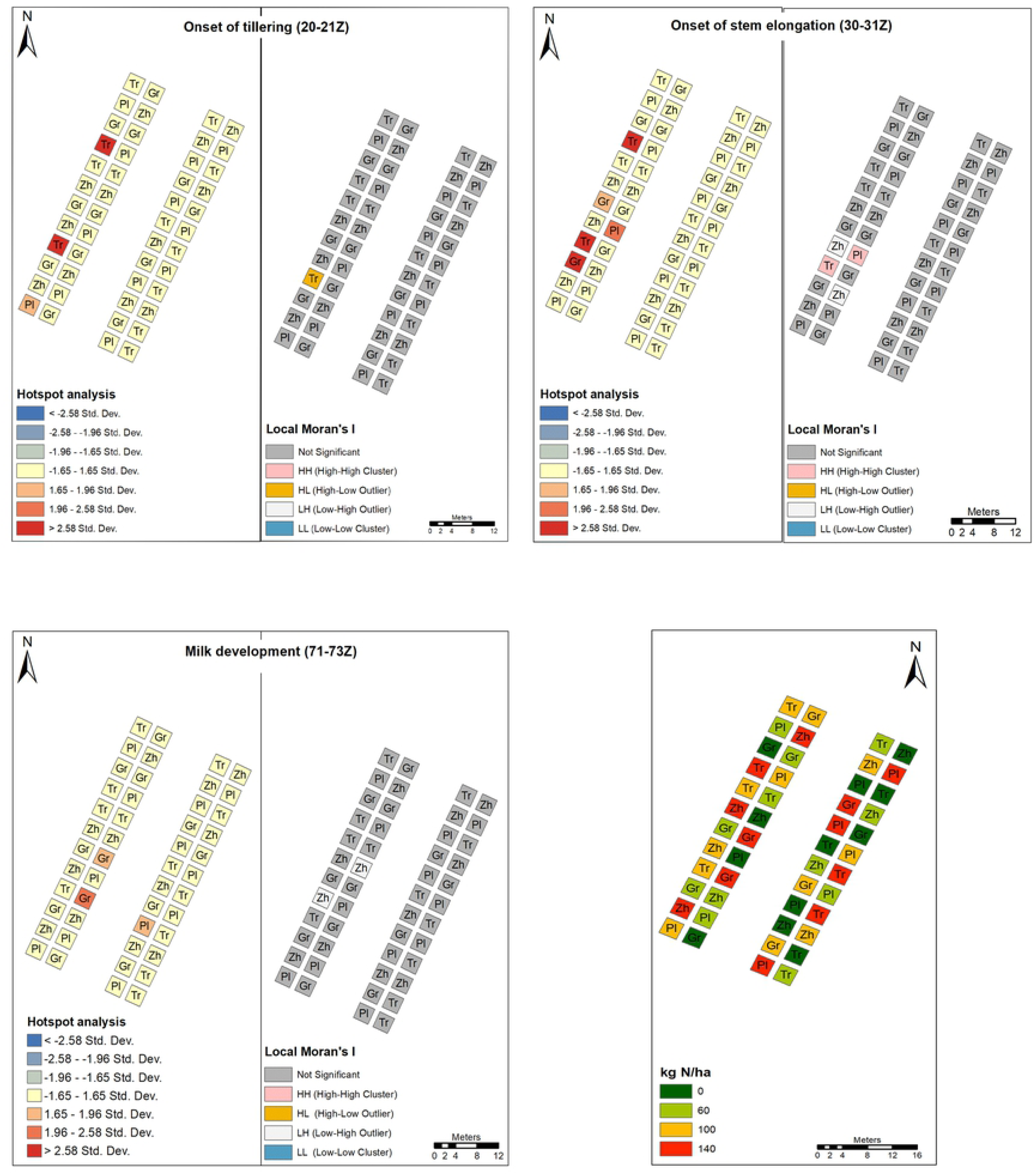
Composite hotspot analysis (Gi z-score) and cluster pattern analysis (Local Moran’s I) of disease severity (caused by *Pyrenophora teres f. teres* and *Rhynchosporium secalis*) assessed at different developmental stages of malt barley. A georeferenced arrangement of the experimental area showing the distribution of the cultivar and N-fertilizer treatments is also presented. The abbreviations stand for: Gr=Grace; Zh=Zhana; Tr=Traveler; Pl=Planet.

**Fig 8.**
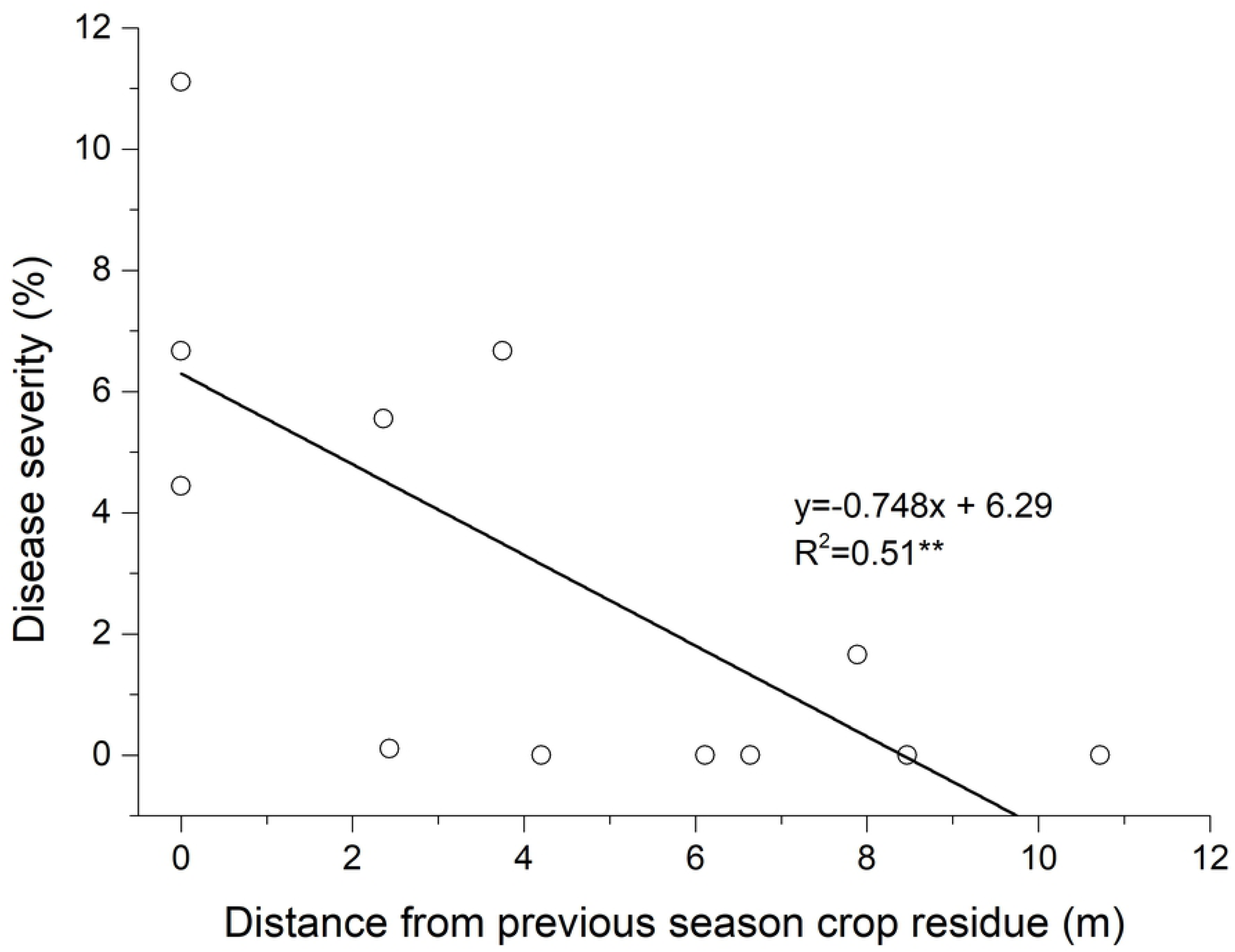
Relationship between disease severity and the distance of Zhana plots from the previous season Zhana’s crop. *At *P*≤ 0.05; **At *P*≤ 0.01; ***At *P*≤ 0.001; ns=non-significant.

During the stem elongation phase, hotspots increased in number and continued to be present in the western part of the field. The analysis identified three hotspots with very high z scores (Grace with 60 kg N /ha; Traveler with 100 kg N /ha; Traveler with 140 kg N /ha, one with high (RGT Planet with 0 kg N /ha) and one with moderate z score (Grace with 60 kg N /ha). Although Zhana with 60 and 100 kg N /ha was surrounded by hot spots, presented low values of disease severity. The Local Moran’s I spatial analysis, confirmed the abovementioned results by characterizing these plots as Low-High outliers, indicating low values of disease severity compared to the surrounding plots. The analysis also identified a statistical significant (p<0.05) cluster of increased disease severity, which coincided with two of the hotspots (Traveller and Planet in the western side) determined with Getis-Ord G* statistic (Fig 7).

Two Grace plots with 140 kg N /ha were identified as hot spots of highest z scores during milk development and followed by RGT Planet without nitrogen application. The Local Moran’s I spatial analysis again identified two Zhana plots (i.e. nitrogen rate 0 and 100 kg/ha) as spatial outliers, since they presented low disease severity in a neighborhood of high values (Fig 7).

### Quantifying the effect of the rate of applied nitrogen and the distance from the nearest hotspot on crop disease severity

Commonality analysis (CA) served to quantify the relative contribution of the rate of applied nitrogen (kg/ha) and the distance from the nearest hotspot on crop disease severity. It is a method of partitioning variance which can discriminate the synergistic or antagonistic processes operating among predictors. Commonalities represent the percentage of variance in the dependent variable that is uniquely explained by each predictor (Unique effect) or by all possible combinations of predictors (Common effect) and their sum is always equal to *R*^*2*^ of the multiple linear regression. The distance from the nearest hotspot (m) and the quantity of applied nitrogen (kg/ha) explained from 10 to 74% of the variance in disease severity (Table 2). Examining the unique effects, it was found, that for the period of stem elongation phase, the distance from the nearest hotspot (m) was the best predictor of disease severity for all the study cultivars, uniquely explaining from 16.8 to 45.5 of its variation. This amount of variance represented from 38.76 to 97.65% of the *R*^*2*^ effect (Table 2). On the contrary, during the onset of grain filling phase the variation in disease severity was best explained by either the nitrogen rate (i.e. Traveler and Grace) or the distance from the nearest hotspot (m) (i.e. RGT Planet and Zhana) (Table 2).

**Table 2.** Commonality coefficients including both unique and common effects, along with % total contribution of each predictor variable or sets of predictor variables to the regression effect.

| Cultivar | Unique and Common Effects | Onset of stem elongation |  | Onset of grain filling (milk development) |  |
| --- | --- | --- | --- | --- | --- |
|  |  | Coefficient | % Total | Coefficient | % Total |
| Traveler | Unique to Distance<sup>a</sup> | 0.4547 | 72.51 | 0.0008 | 0.22 |
|  | Unique to Nitrogen<sup>b</sup> | 0.0004 | 0.07 | 0.3493 | 93.17 |
|  | Common to Distance, and Nitrogen | 0.1720 | 27.42 | 0.0248 | 6.61 |
|  | Total | 0.6271 | 100.00 | 0.3748 | 100.00 |
| Zhana | Unique to Distance | 0.1678 | 67.81 | 0.0819 | 79.91 |
|  | Unique to Nitrogen | 0.0089 | 3.59 | 0.0241 | 23.51 |
|  | Common to Distance, and Nitrogen | 0.0708 | 28.61 | -0.0035 | -3.42 |
|  | Total | 0.2475 | 100.00 | 0.1025 | 100.00 |
| Grace | Unique to Distance | 0.3837 | 97.65 | 0.1641 | 22.26 |
|  | Unique to Nitrogen | 0.0105 | 2.66 | 0.2850 | 38.66 |
|  | Common to Distance, and Nitrogen | -0.0012 | -0.31 | 0.2881 | 39.08 |
|  | Total | 0.3930 | 100.00 | 0.7373 | 100.00 |
| RGT Planet | Unique to Distance | 0.1912 | 38.76 | 0.3672 | 83.26 |
|  | Unique to Nitrogen | 0.0925 | 18.75 | 0.0020 | 0.46 |
|  | Common to Distance, and Nitrogen | 0.2096 | 42.49 | 0.0718 | 16.29 |
|  | Total | 0.4933 | 100.00 | 0.4411 | 100.00 |

<sup>a</sup>Refers to the distance from the nearest hotspot (m)
<sup>b</sup>Refers to the rate of applied nitrogen (kg/ha)

## Discussion

Our results demonstrated that both NFNB and leaf scald can be carried over from one season to the next on infected seed under Mediterranean conditions, which is in line with previous reports [3, 39]. Typical yield losses due to NFNB (*Pyrenophora teres f. teres*) and leaf scald (*Rhynchosporium secalis*) outbreaks can be up to 30–40% [3, 6, 8-11]. However, we did not detect any consistent relationship between disease severity and grain yield when the main source of variation was nitrogen rate (Fig 6). Jalakas et al. [40] also found a weak relationship between malt barley grain yield and net blotch (*Pyrenophora teres*) disease severity. This can be attributed to the time of disease occurrence and the extent of disease severity in relation to barley developmental stage. It is widely accepted that grain yield determination in barley is mainly explained by the variation of grain number per unit of land area [20, 33, 41-42]. According to Bingham et al. [43], grain number in barley is a function of the production and survival of tillers and spikelets and the success of fertilization of florets. Tiller production and spikelet initiation occur before stem elongation phase, while the survival and further growth of tillers and spikelets is largely determined from stem elongation onwards. Accordingly, our results showed that the highest disease severity, which was recorded in Traveler during tillering phase (Fig 2), exerted a more pronounced negative effect on grain yield (Fig 6). In addition, the higher disease severity in Grace compared to the rest of the studied cultivars during the onset of grain filling phase (Fig 2), led to a significant reduction in grain yield, mainly through a decrease in mean grain weight. Indeed, an increase in disease severity by 32.5% during grain filling phase caused a reduction in thousand grain weight by 18.3% in Grace. In line with this, Agostinetto et al. [44] demonstrated that the strongest relationship between grain yield reduction and barley spot blotch severity occurred after the booting stage of barley. Furthermore, Khan [9] observed a reduction in barley grain yield by 25-35% from net blotch, mainly due to a significant decrease in thousand grain weight.

The effect of N on plant disease severity is quite variable in literature [27]. Both increases [13, 25, 28] and decreases [26] of disease severity are reported by increasing N in plants. In addition, Turkington et al. [12] found that total leaf disease severity, caused by NFNB, in barley was not significantly affected by N rate. Our results showed that disease severity for both pathogens tended to increase from anthesis onwards by increasing the rate of applied nitrogen (Fig 4). This relationship can be attributed to some extent to the synergistic effect of the N fertilizer type used in this study. It is suggested that nitrate fertilizers increase the severity of disease whereas ammonium fertilizers decrease it [28 and references quoted therein].

Grain protein content is one of the most important factors in marketing malting barley. The primar objective, particularly in Mediterranean environments, is to maintain grain protein content below a threshold of 11.5-12.0% depending on brewing industry [33]. Although there is some evidence from northern climates suggesting that NFNB infections are not exerting any significant effect on grain protein content [12-13], our results revealed for the first time a positive relationship between NFNB disease severity and grain protein content under Mediterranean conditions. Additionally, it was shown that the magnitude of this relationship was genotype dependent (Fig 5). It seems that the effect of NFNB disease severity on grain protein content increases under terminal drought stress conditions in April-May (Figs 1A and 1B). According to Bertholdsson [17] drought stress during late grain filling, limits carbohydrate incorporation in the grain and causes pre-maturation and less dilution of the protein in the grain.

The epidemiology assessment of both diseases, when nitrogen rate and genotype were the main sources of variation, was implemented with hotspot and Anselin Local Moran’s I analysis. The location of hotspots was modified during the growing season (Fig 7). This can be explained either by the soil heterogeneity or by the spatial presence of the pathogens in the soil (i.e. as infected host residue) and genotype susceptibility. The soil heterogeneity was considered negligible because: i) the acreage of the experimental field was small (approximately 0.1 ha), ii) there was no land inclination and iii) the differentiation of the field soil moisture was rather small (Fig 9).

**Fig 9.**
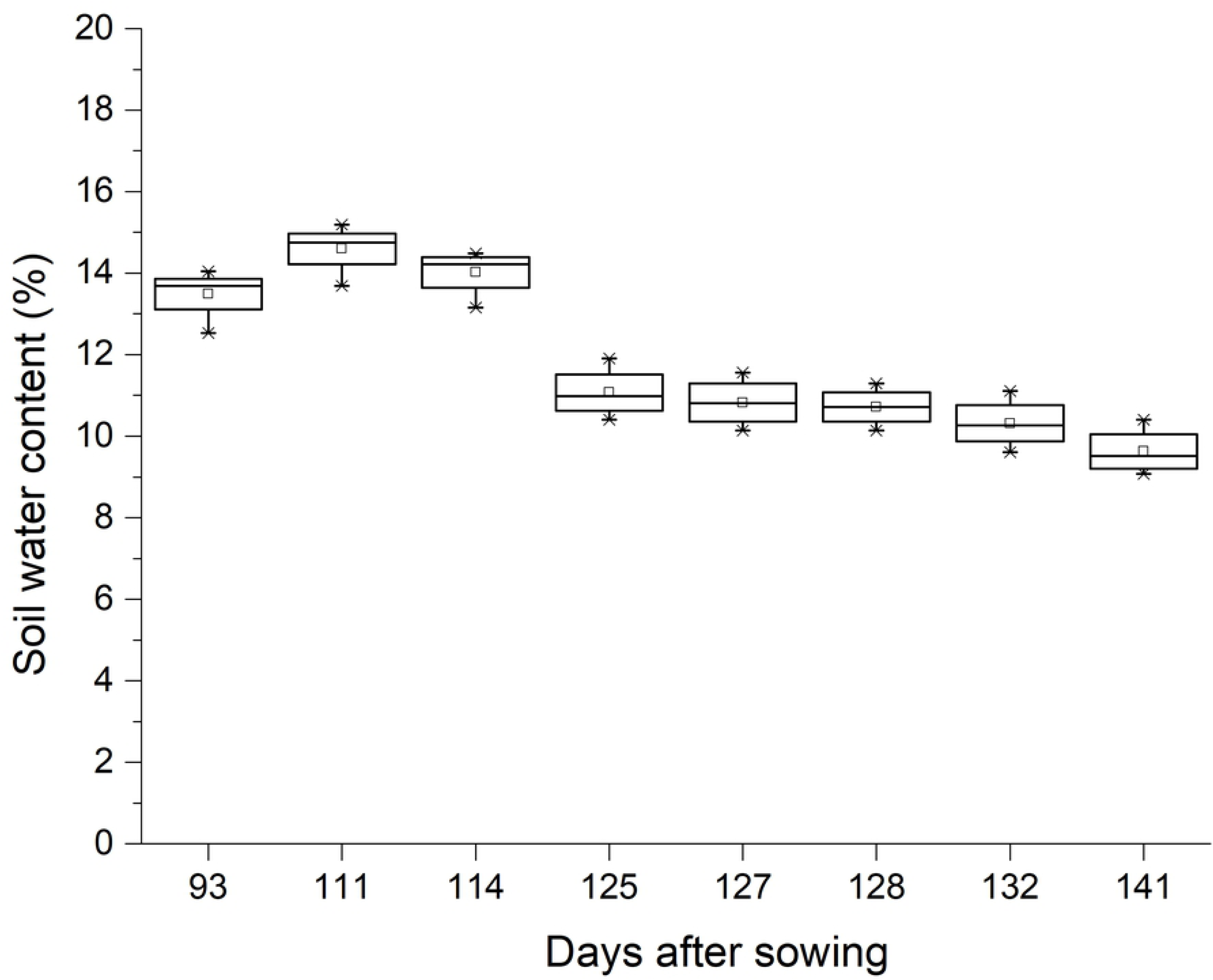
The variation in soil water content from anthesis until the end of grain filling (during Exp2). Broad lines are medians, square open dots are means, boxes show the interquartile range and whiskers extend to the last data point within 1.5 times the inter-quartile range.

Commonality analysis, when nitrogen rate and genotype were the main sources of variation, revealed that the most important factor concerning NFNB disease severity was the distance of plots from the hotspots, concerning the period of onset of stem elongation (Table 2). According to Liu et al. [4], NFNB is classified as stubble-borne disease because the fungus usually produces the ascocarp as an over-seasoning structure on infected barley debris left after harvest. The primary inoculum early in the growing season is made by mature ascospores which are dispersed by wind. After initial colonization, the pathogen produces a large number of conidia which serve as secondary inocula. These asexually produced spores can be dispersed either by wind or rain to cause new infections on plants locally or at longer distances [4 and references quoted therein].

On the other hand, Zhana was the only cultivar which was not infected by NFNB during neither seasons (i.e. it was infected only by *Rhynchosporium secalis*). However, it was found that the distance of Zhana experimental plots from the previous season crop residues (i.e. the sites with Zhana) explained 51% of the variation in disease severity (Fig 8). This result is also supported by the Anselin Local Moran’s I spatial statistical analysis. Zhana was considered an outlier due to lower disease severity values although surrounded by plots with high values from stem elongation onwards (Fig 7).

The late occurrence of *Rhynchosporium secalis* symptoms on Zhana compared to NFNB (Fig 2) during both experiments, can be possibly attributed to its specific life cycle. According to Zhan et al. [3] *R. secalis* grows symptomlessly under the cuticle, especially where walls of adjacent cells are joined before producing new conidia and finally, visual symptoms. Further investigations concerning the infection process of *R. secalis* in barley had been conducted by Linsell et al. [45].

In general, NFNB was more prevalent compared to leaf scald during all tested developmental phases of malt barley (Figs 2 and 3). According to Robinson and Jalli [46] this could be a result of net blotch being comparatively less demanding of environmental conditions (mostly wind dispersed) than scald (mostly splash dispersed) for effective spore dispersal and epidemic development.

## Conclusions

The results of the present study provide a further insight into the epidemiology and the effect of nitrogen fertilization on the most important foliar diseases of malt barley in Greece. It was demonstrated that both NFNB and leaf scald can be carried over from one season to the next on infected seed under Mediterranean conditions. However, disease severity was more pronounced after barley tillering phase when soil had been successfully enriched first with the pathogen propagules. When both plant pathogens were present in soil residues, it was shown that the effect of the distance of cultivars from hotspots (i.e. the locations with the highest disease infections) was a better predictor of disease severity (for both diseases) compared to nitrogen rate during the pre-anthesis period. However, after anthesis disease severity was best explained by nitrogen rate concerning the most susceptible cultivars to NFNB. In addition, it was presented that the effect of disease infections on yield, grain size and grain protein content varied in relation to genotype, pathogen and the stage of crop development.

## Acknowledgements

The present study was funded by the Athenian Brewery SA through the project “Evaluation of macro and micro barley trials of Athenian Brewery SA”.

## References

1. Friedt W. Barley breeding history, progress, objectives, and technology. In: Barley production, improvement, and uses. Ullrich, S.E. (ed), Wiley-Blackwell. 2011; pp. 160–220

2. Meussdoerffer F. and Zarnkow M. Starchy Raw Materials. In: Handbook of brewing: processes, technology, markets. Eßlinger, H.M. (ed), Wiley-VCH Verlag GmbH & Co. KGaA, Weinheim. 2009; pp. 43–83

3. Zhan J, Fitt BDL, Pinnschmidt HO, Oxley SJP, Newton AC. Resistance, epidemiology and sustainable management of Rhynchosporium secalis populations on barley. Plant Pathol. 2008; 57, 1–14.

4. Liu Z, Ellwood SR, Oliver RP, Friesen TL. Pyrenophora teres: profile of an increasingly damaging barley pathogen. Mol Plant Pathol. 2011; 12, 1–19.

5. Paulitz TC and Steffenson BJ. Biotic stress in barley: Disease problems and solutions. In: Barley production, improvement, and uses. Ullrich, S.E. (ed), Wiley-Blackwell. 2011. pp. 307–354

6. Shipton WA. Effect of net blotch infection of barley on grain yield and quality. Austral J Exp Agr and Animal Hus. 1966; 6: 437–440.

7. Shipton WA, Boyd WJR and Ali SM. Scald of barley. Reviews in Plant Patholology 1974; 53: 839–861.

8. Martin RA. Disease progression and yield loss in barley associated with net blotch, as influenced by fungicide seed treatment. Can J Plant Pathol 1985; 7: 83–90.

9. Khan TN. Relationship between net blotch (Drechslera teres) and losses in grain yield of barley in Western Australia. Aust J Agr Res. 1987; 38: 671–679.

10. El Yousfi B. and Ezzahiri B. Net blotch in semi-arid regions of Marocco II: Yield and yield-loss modeling. Field Crop Res. 2002; 73: 81–93.

11. Murray GM. and Brennan JP. Estimating disease losses to the Australian barley industry. Australas Plant Path. 2010; 39: 85–96.

12. Turkington TK, O’Donovan JT, Edney MJ, Juskiw PE, McKenzie RH, Harker KN, et al. Effect of crop residue, nitrogen rate and fungicide application on malting barley productivity, quality, and foliar disease severity. Can J Plant Sci. 2012; 92: 577–588.

13. Kangor T, Sooväli P, Tamm Y, Tamm I and Koppel M. Malting barley diseases, yield and quality - responses to using various agro-technology regimes. Proc Latvian Academy of Sciences. Section B, Vol. 71 No. 1/2 (706/707). 2017. pp. 57–62.

14. Grashoff, C and D’Antuono LF. Effect of shading and nitrogen application on yield, grain size distribution and concentrations of nitrogen and water soluble carbohydrates in malting spring barley (Hordeum vulgare L.). Eur J Agron. 1997; 6: 275–293.

15. Bingham IJ, Blake J, Foulkes MJ and Spink J. Is barley yield in the UK sink limited? II. Factors affecting potential grain size. Field Crop Res. 2007; 101: 212–220.

16. Ugarte C, Calderini DF and Slafer GA. Grain weight and grain number responsiveness to pre-anthesis temperature in wheat barley and triticale. Field Crop Res. 2007; 100: 240–248.

17. Bertholdsson, NO. Characterization of malting barley cultivars with more or less stable grain protein content under varying environmental conditions. Eur J Agron. 1999; 10: 1–8.

18. Baethgen WE, Christianson CB and Lamothe AG. Nitrogen fertilizer effects on growth, grain yield, and yield components of malting barley. Field Crop Res. 1995; 43: 87–99.

19. Abeledo LG, Calderini DF and Slafer GA. Nitrogen economy in old and modern malting barleys. Field Crop Res. 2008; 106: 171–178.

20. Albrizio R, Todorovic M, Matic T and Stellacci AM, Comparing the interactive effects of water and nitrogen on durum wheat and barley grown in a Mediterranean environment. Field Crop Res. 2010; 115: 179–190.

21. Dordas C. Variation in dry matter and nitrogen accumulation and remobilization in barley as affected by fertilization, cultivar, and source–sink relations. Eur J Agron. 2012; 37: 31–42.

22. Grant CA, Gauer LE, Gehl DT, and Bailey LD. Protein production and nitrogen utilization by barley cultivars in response to nitrogen fertilizer under varying moisture conditions. Can J Plant Sci. 1991; 7l: 997–1009.

23. Boonchoo S, Fukai S and Hetherington SE. Barley yield and grain protein concentration as affected by assimilate and nitrogen availability. Aust J Agr Res. 1998; 49: 695–706.

24. Agegnehu G, Nelson PN and Bird MI. The effects of biochar, compost and their mixture and nitrogen fertilizer on yield and nitrogen use efficiency of barley grown on a Nitisol in the highlands of Ethiopia. Sci Total Environ. 2016; 869–879

25. Jenkyn JF and Griffiths E. Relationships between the severity of leaf blotch (Rhynchosporium secalis) and the water-soluble carbohydrate and nitrogen contents of barley plants. Ann Appl Biol. 1978; 90: 35–44.

26. Krupinsky JM, Halvorson AD, Tanaka DL and Merrill SD. Nitrogen and tillage effects on wheat leaf spot diseases in the northern great plains. Agron J. 2007; 99: 562–569.

27. Dordas C. Role of nutrients in controlling plant diseases in sustainable agriculture. A review. Agron Sustain Dev. 2008; 28: 33–46.

28. Veresoglou SD, Barto EK, Menexes G and Rillig MC. Fertilization affects severity of disease caused by fungal plant pathogens. Plant Pathol. 2013; 62: 961–969.

29. Zadoks JC, Chang TT and Konzak CF. A decimal code for the growth stages of cereals. Weed Res. 1974; 14: 415–421.

30. Saari EE and Prescott JM. Scale for appraising the foliar intensity of wheat diseases. Plant Dis Rep. 1975; 59: 377–380.

31. Shaner G and Finney R. The effect of nitrogen fertilization on the expression of slow-mildewing resistance in knox wheat. Phytopathology. 1977; 67: 1051–1056.

32. Analytica EBC. Sieving Test for Barley Method 3.11. 1998.

33. Vahamidis P, Stefopoulou A, Kotoulas V, Lyra D, Dercas N and Economou G, Yield, grain size, protein content and water use efficiency of null-LOX malt barley in a semiarid Mediterranean agroecosystem. Field Crop Res. 2017; 206; 115–127.

34. Getis, A and Ord JK. The analysis of spatial association by use of distance statistics. Geogr Anal. 1992; 24: 189–206.

35. Zhang H and Tripathi NK. Geospatial hot spot analysis of lung cancer patients correlated to fine particulate matter (PM2.5) and industrial wind in Eastern Thailand. J Clean Prod. 2018; 170: 407–424.

36. Zhang C, Luo L, Xu W and Ledwith V. Use of local Moran’s I and GIS to identify pollution hotspots of Pb in urban soils of Galway, Ireland. Sci Total Environ. 2008; 398: 212–221.

37. Lalor GC and Zhang C. Multivariate outlier detection and remediation in geochemical databases. Sci Total Environ. 2001; 281: 99–109.

38. Nimon K, Lewis M, Kane R and Haynes RM. An R package to compute commonality coefficients in the multiple regression case: An introduction to the package and a practical example. Behav ResMethods. 2008; 40: 457–466.

39. McLean MS, Howlett BJ and Hollaway GJ. Epidemiology and control of spot form of net blotch (Pyrenophora teres f. maculata) of barley: a review. Crop Pasture Sci. 2009; 60: 303–315.

40. Jalakas P, Tulva I, Kangor T, Sooväli P, Rasulov B, Tamm Ü et al. Gas exchange-yield relationships of malting barley genotypes treated with fungicides and biostimulants. Eur J Agron. 2018; 99: 129–137.

41. Cossani CM, Savin R and Slafer GA. Contrasting performance of barley and wheat in a wide range of conditions in Mediterranean Catalonia (Spain). Ann Appl Biol. 2007; 151: 167–173.

42. Cossani CM, Slafer GA and Savin R. Yield and biomass in wheat and barley under a range of conditions in a Mediterranean site. Field Crop Res. 2009; 112: 205–213.

43. Bingham IJ, Hoad SP, Thomas WTB and Newton AC. Yield response to fungicide of spring barley genotypes differing in disease susceptibility and canopy structure. Field Crop Res. 2012; 139: 9–19.

44. Agostinetto L, Casa RT, Bogo A, Sachs C, Souza CA, Reis EM et al. Barley spot blotch intensity, damage, and control response to foliar fungicide application in southern Brazil. Crop Prot. 2015; 67: 7–12.

45. Linsell KJ, Keiper FJ, Forgan A and Oldach KH. New insights into the infection process of Rhynchosporium secalis in barley using GFP. Fungal Genet Biol. 2011; 48: 124–131.

46. Robinson J and Jalli M. Grain yield, net blotch and scald of barley in Finnish official variety trials. Agr Food Sci Finland. 1997; 6: 399–408.

